# *APOL1-G0* protects podocytes in a mouse model of HIV-associated nephropathy

**DOI:** 10.1101/598557

**Authors:** Leslie A. Bruggeman, Zhenzhen Wu, Liping Luo, Sethu Madhavan, Paul E. Drawz, David B. Thomas, Laura Barisoni, John F. O’Toole, John R. Sedor

**Affiliations:** Departments of Inflammation & Immunity, Case Western Reserve University School of Medicine, Case Western Reserve University, Cleveland, OH; Departments of Nephrology, Cleveland Clinic Lerner College of Medicine, Case Western Reserve University School of Medicine, Case Western Reserve University, Cleveland, OH; Departments of Physiology and Biophysics, Case Western Reserve University School of Medicine, Case Western Reserve University, Cleveland, OH, and Department of Medicine, University of Minnesota, Minneapolis, MN; Departments of Pathology, University of Miami, Miami, FL; Departments of Pathology and Medicine, Division of Nephrology, Duke University, Durham, NC

## Abstract

**Background:** African polymorphisms in the gene for Apolipoprotein L1 (*APOL1*) confer a survival advantage against lethal trypanosomiasis but also an increased risk for several chronic kidney diseases (CKD) including HIV-associated nephropathy (HIVAN). APOL1 is expressed in renal cells, however, the pathogenic events that lead to renal cell damage and kidney disease are not fully understood.

**Methods:** The podocyte function of *APOL1-G0* versus *APOL1-G2* in the setting of a known disease stressor was assessed using transgenic mouse models. Survival, renal pathology and function, and podocyte density were assessed in an intercross of a mouse model of HIVAN (Tg26) with two mouse models that express either *APOL1-G0* or *APOL1-G2* in podocytes.

**Results:** Mice that expressed HIV genes developed heavy proteinuria and glomerulosclerosis, and had significant losses in podocyte numbers and reductions in podocyte densities. Mice that co-expressed *APOL1-G0* and HIV had preserved podocyte numbers and densities, with fewer morphologic manifestations typical of HIVAN pathology. Podocyte losses and pathology in mice co-expressing *APOL1-G2* and HIV were not significantly different from mice expressing only HIV. Podocyte hypertrophy, a known compensatory event to stress, was increased in the mice co-expressing HIV and *APOL1-G0*, but absent in the mice co-expressing HIV and *APOL1-G2*. Mortality and renal function tests were not significantly different between groups.

**Conclusions:** *APOL1-G0* expressed in podocytes may have a protective function against podocyte loss or injury when exposed to an environmental stressor. This function appears to be absent with *APOL1-G2* expression, suggesting *APOL1-G2* is a loss-of-function variant.

## INTRODUCTION

Polymorphisms in the gene for Apolipoprotein L1 (APOL1, gene name *APOL1*) are found only in populations of recent African ancestry and confer significant risk for chronic kidney diseases (CKD) including HIV-associated nephropathy (HIVAN), idiopathic focal segmental glomerulosclerosis (FSGS), and hypertension-attributed CKD.^1-5^ APOL1 is constitutively secreted into the blood and functions to kill trypanosome parasites that cause African sleeping sickness. The CKD-associated risk alleles, known as variants G1 and G2, kill a broader range of parasites compared to the common allele (known as G0) and provide an evolutionary survival advantage (reviewed in^6^). A single variant *APOL1* allele is sufficient to protect against trypanosomiasis however, risk for kidney disease is recessive requiring two variant alleles. Many individuals with a high risk genotype of two *APOL1* variants, however, do not develop kidney disease. Thus, APOL1-associated CKDs appear to be a gene-environment dependent process where the genetic susceptibility manifests in disease only when the individual is exposed to a triggering environmental stimulus.

Although the trypanolytic APOL1 in blood is abundant, studies to date have not associated circulating APOL1 with CKD risk, an observation corroborated by poorer kidney transplant outcomes dependent on donor *APOL1* genotype.^7-11^ *APOL1* is also expressed in some renal cells including the podocyte.^12-14^ In HIVAN and mouse models of HIVAN, HIV-1 genes also are expressed in podocytes,^15-22^ and HIV-1 gene expression in podocytes alone is sufficient to be disease-causing in mouse models.^23, 24^ Thus, HIVAN is an ideal disease to study the functional interaction of podocyte-expressed *APOL1* with a known environmental trigger (HIV).

An intercross between APOL1 transgenic mice with a mouse model of HIVAN would provide an *in vivo* system to examine the podocyte function of *APOL1-G0* and *APOL1-G2* in the setting of a known human disease stressor. Predictions were either disease exacerbation if the *APOL1* variants contribute a deleterious function, or alternatively, disease mitigation if *APOL1-G0* provides a beneficial function. After assessment of renal function and pathology, *APOL1-G2* did not exacerbate the HIVAN phenotype. *APOL1-G0*, however, reduced podocyte losses and glomerulosclerosis suggesting *APOL1-G0* provided some protection against glomerular injury caused by HIV.

## METHODS

### Mouse models and phenotyping

All animal studies adhered to the *NIH Guide for the Care and Use of Laboratory Animals* and were conducted under oversight by Case Western Reserve University. The transgenic mouse models for podocyte-restricted expression of human *APOL1-G0* (“Tg-G0” F38 line) and *APOL1- G2* (“Tg-G2” F24 line) using the *Nphs1* promoter have matched glomerular expression patterns and were previously described.^25^ The Tg26/*HIVAN4* mouse model of HIVAN is a congenic of Tg26^26^ that develops less severe kidney disease and has been previously described.^27^ The *APOL1* transgenic are on the FVB/N background and the Tg26/*HIVAN4* model is >99% FVB/N with a 60Mb BALB/c-derived genomic region referred to as the *HIVAN4* locus.^27^ The Tg26/*HIVAN4* and *APOL1* transgenic models are maintained as carriers (hemizygotes), thus the intercross generated all possible single and dual transgenics for age-matched comparisons. Mice were housed in a specific pathogen free conventional animal facility and standard breeding practices were used to generate F1 hybrids resulting in the expected Mendelian proportions of: 25% non-transgenic (wildtype), 25% *APOL1* single transgenic, 25% Tg26/*HIVAN4* single transgenic, and 25% *APOL1* plus Tg26/*HIVAN4* dual transgenic mice. All dual transgenics carried a single copy of the respective *APOL1* gene similar to the single transgenics.

Two hundred day-old F1 hybrids (n=18-21 each group, combined males and females) were phenotyped for kidney disease typical of HIVAN. Renal function testing was performed by the Vanderbilt Center for Kidney Disease Pathology and Phenotyping Core and included ELISAs for urinary albumin and creatinine and HPLC assays for serum creatinine. Kidneys were PAS stained and glomerular and tubular pathology was scored using quantitative methods by pathologists blinded to sample identity. Total number of glomeruli were counted, and percentages of glomeruli with the following features were calculated: segmental sclerosis, global sclerosis, segmental collapse, global collapse, podocyte hypertrophy (of ≥1 podocyte with large/prominent cell body with or without increased size of the nucleus), podocyte hyperplasia (of ≥2 layers of normal or hypertrophic podocytes). Tubular microcysts (dilated tubule, often with serpiginous appearance, containing a large hyaline cast), tubular atrophy, interstitial fibrosis, and interstitial inflammation also were scored using a semi-quantitative scale of 0 to 4, where 0= unaffected; 0.5= 1-5% affected; 1= 6-25% affected; 2= 26-50% affected; 3= 51-75% affected; 4= >75% affected.

### Imaging and podocyte density

Kidney sections were examined using immunofluorescence and confocal microscopy as described previously.^14^ Primary antibodies used were rabbit polyclonal APOL1 (1:400 dilution, Sigma), mouse monoclonal Synaptopodin (1:10 dilution, BioDesign), rabbit polyclonal WT-1 (1:200 dilution, Santa Cruz Biotech). Podocyte counts, glomerular volumes, and podocyte density calculations used a method originally described by Venkatareddy *et al*.^36^ and as used previously for characterization of the Tg-G0 and Tg-G2 mouse models.^25^

### Statistical methods

Podocyte density calculations were based on ∼50 glomeruli for each animal with combined male and female mice per group (actual group numbers are in figure legends). Group differences were analyzed using ANOVA. Generalized linear mixed models and Markov Chain Monte Carlo samples were used to evaluate differences in podocyte density, glomerular volume, and corrected podocyte count between groups. Differences in renal function tests and histopathology scoring between groups were determined by *t* test followed by Bonferroni correction. *P* values ≤0.05 were considered significant. Animals in the phenotyping study that died or were euthanized for humane reasons prior to the 200 day endpoint were included in the analysis. Inclusion of these animals did not alter study outcomes by sensitivity analyses.

## RESULTS

Two transgenic mouse models expressing either the human *APOL1-G0* (“Tg-G0”) or *APOL1-G2* (“Tg-G2”) genes under control of the Nephrin (*Nphs1*) promoter, restrict *APOL1* expression to podocytes, but do not spontaneously develop CKD.^25^ Since mice do not have an ortholog of human *APOL1*, the Tg26/*HIVAN4* mice would represent an *APOL1* null phenotype, and mice expressing *APOL1-G0* would functionally recreate humans homozygous for *APOL1-G0* (a CKD low risk genotype) whereas mice expressing *APOL1-G2* would functionally recreate humans homozygous for *APOL1-G2* (a CKD high risk genotype). The Tg26/*HIVAN4* mouse model of HIVAN is transgenic for a subgenomic HIV-1 provirus and spontaneously develops a progressive and lethal kidney disease that replicates most of the pathology and clinical presentation of the human disease.^26, 27^ F1 hybrids from intercrossing Tg26/*HIVAN4* with Tg-G0 or Tg-G2 (dual transgenics referred to as “Tg26+G0” and “Tg26+G2”) were examined at 200 days of age for pathology and renal function using parameters previously established to quantitate disease severity in the Tg26 model.^28^ Age-matched non-transgenic (wildtype), Tg26/*HIVAN4*, and Tg-G0 and Tg-G2 single transgenics were also examined as comparators.

### Renal pathology

Two pathologists independently assessed (scored) ten different parameters of glomerular and tubulointerstitial damage (**Supplemental Table 1**). An overall composite disease score integrating all categories was not significantly different between groups, although the Tg26+G0 mice trended toward less severe pathology compared to Tg26/*HIVAN4* and Tg26+G2 (**Table 1**). Several individual glomerular features, however, were different. The percentage of sclerotic glomeruli trended lower in the Tg26+G0 hybrid mice (**Figure 1A**). In addition, podocyte hypertrophy was greater in Tg26+G0 mice compared to Tg26+G2 mice (**Figure 1B and Supplemental Table 1**). Only one Tg26+G2 mouse exhibited sclerotic glomeruli with hypertrophic podocytes, whereas hypertrophy was evident in diseased glomeruli of over half of the Tg26+G0 mice. All tubulointerstitial features were not different between groups (**Supplemental Table 1**).

**Table 1:** Renal function in single and dual transgenic mice.

|  | **n**<br>**(%male)** | **Serum Creatinine/**<br>**body weight**<br>**(mg/dl/kg)**<br>**mean ± SD** | ***P*** | **UACR**<br>**(μg/mg)**<br>**median (IQR)** | ***P*** | **Composite**<br>**Histopathology**<br>**Score**<br>**median (IQR)** | ***P*** |
| --- | --- | --- | --- | --- | --- | --- | --- |
| **Tg26** | 21 (71%) | 4.2 ± 0.9 | ref | 212 (84,493) | ref | 0.36 (0.14, 0.43) | ref |
| **Tg26+G0** | 21 (62%) | 6.8 ± 5.2 | NS | 141 (108, 226) | NS | 0.21 (0.14, 0.71) | NS |
| **Tg26+G2** | 18 (33%) | 4.2 ± 1.3 | NS | 119 (97, 179) | NS | 0.25 (0.07, 0.50) | NS |

UACR, urinary albumin to creatinine ratio
SD, standard deviation
IQR, interquartile range
NS, not significant

**Figure 1.**
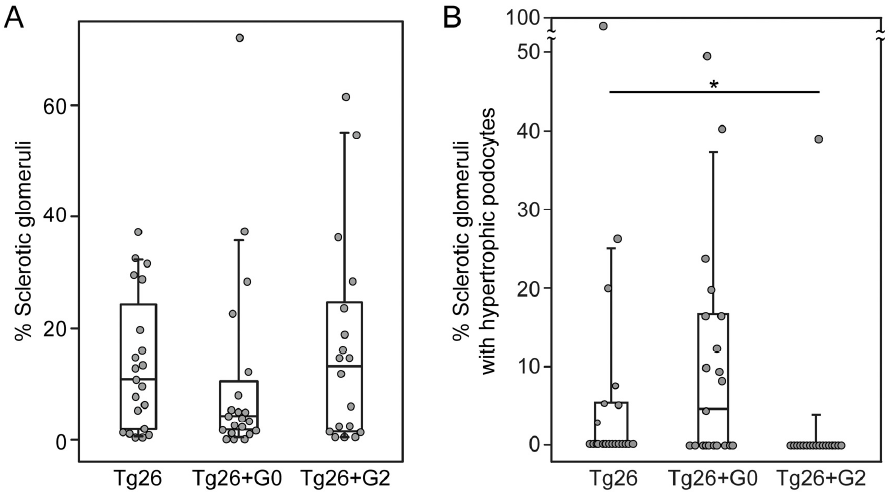
*APOL1-G0* reduced glomerulosclerosis in a murine model of HIVAN. Box whisker plots of the significantly different pathological features from the histopathologic disease scoring presented in Supplemental Table 1. **A**. Total sclerotic glomeruli (aggregate global and sclerotic glomeruli from supplemental table 1) as a percentage of total scored glomeruli per animal. Differences were not significant, but with exclusion of the outlier (#) in the Tg26+G0 group, this group approached significance (*P*=0.08) compared to both Tg26 and Tg26+G2. **B**. Percentage of sclerotic glomeruli with hypertrophic podocytes, only one mouse had evidence of podocyte hypertrophy in the Tg26+G2 group. Each data point represents one mouse, boxes are interquartile range, whiskers are 95% confidence intervals. Numbers in each group were Tg26 n=21, Tg26+G0 n=21, Tg26+G2 n=18. Statistical comparisons were made to the non-*APOL1* expressing group (Tg26+G0 or Tg26+G2 versus Tg26;\**P*<0.05).

### Podocyte density

The podocyte depletion hypothesis purports chronic glomerular diseases are mediated by progressive podocyte losses via podocyte detachment or death. It has been validated in many human and rodent disease models including the age-related decline in renal function (reviewed in^29^). Although the Tg-G0 and Tg-G2 transgenic mice do not spontaneously develop kidney disease, the Tg-G2 mice have an accelerated age-related decrease in podocyte densities compared to Tg-G0 or wild-type mice.^25^ This accelerated podocyte loss is not associated with podocyte cell death and remained subclinical, with longitudinal predictions that podocyte attrition would remain insufficient to initiate glomerulosclerosis through the average mouse lifespan.^30^ It is unknown whether this accelerated podocyte depletion could be exacerbated by a disease stressor, and thus, underlie a pathogenic mechanism of the *APOL1* variants.

In Tg26/*HIVAN4* mice, podocyte densities were significantly less compared to all other non-diseased groups as would be expected for the progressive glomerular disease that occurs in the model at 200 days of age. Podocytes were lost segmentally, exhibiting losses in WT-1 positivity (podocyte number) but with preserved Synaptopodin staining (glomerular volume), a finding that reflects possible compensatory hypertrophy by the residual podocytes. More severely affected glomeruli exhibited losses in both WT-1 positivity and Synaptopodin staining, resulting in reductions in both podocyte number and glomerular volume which occurs when podocyte loss exceeds the adaptive capacity of the remaining podocytes. (**Supplemental Figure 1**). Podocyte densities in the dual transgenic Tg26+G0 and Tg26+G2 were both significantly reduced compared to the single transgenic Tg-G0 and Tg-G2 mice due to the glomerulosclerosis of the Tg26/*HIVAN4* model (**Figure 2A**). Podocyte densities in the Tg26/*HIVAN4* and Tg26+G2 dual transgenic mice were not significantly different. However, podocyte densities in the Tg26+G0 dual transgenics were significantly greater than either the Tg26+G2 dual transgenic and Tg26/*HIVAN4* mice. This preservation of podocyte density was driven by higher numbers of podocytes (**Figure 2B**) since glomerular volumes were not significantly different (**Supplemental Figure 2**). This suggests podocyte *APOL1-G0* expression functioned to reduce podocyte loss in the setting of HIVAN-like kidney disease.

**Figure 2.**
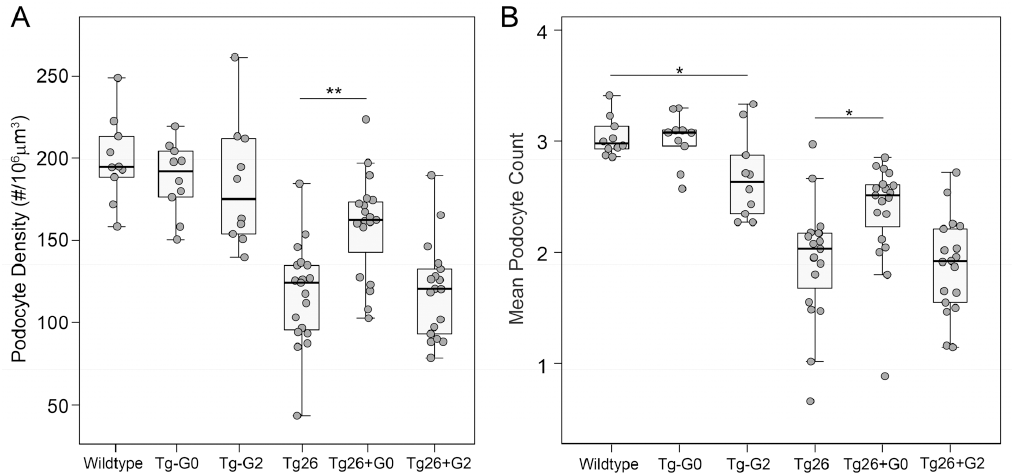
*APOL1-G0* preserves podocyte density in a murine model of HIVAN. **A**. Mean podocyte densities calculated from podocyte counts (panel B) and glomerular volumes (shown in Supplemental Figure 2). **B**. Mean podocyte number per glomeruli. Each data point represents one mouse, boxes are interquartile range, whiskers are 95% confidence intervals. Numbers in each group were: Non-transgenic (wild-type) n=10, Tg26 n=19, Tg26+G0 n=19, Tg26+G2 n=18, Tg-G0 n=10, Tg-G2 n=10. Statistical comparisons were made to the relevant non-*APOL1* expressing group (Tg-G0 or Tg-G2 versus wildtype, and Tg26+G0 or Tg26+G2 versus Tg26; \**P*<0.05, \*\**P*<0.01).

### Renal function

Standard renal function tests for serum creatinine and urinary albumin to creatinine ratios were not significantly different in any of the groups (**Table 1**). As expected, some animals died or reached predetermined humane endpoints and were sacrificed before study end, but there was no significant difference in survival in any group (**Supplemental Figure 3**).

**Figure 3.**
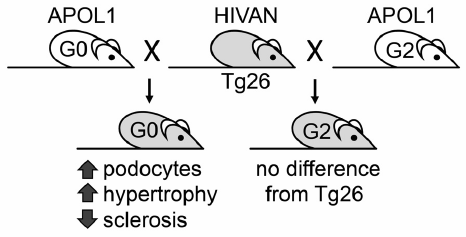
Podocyte *APOL1-G2* expression replicates *APOL1* null phenotype suggesting the *G2* polymorphism is a loss-of-function mutation. Characterization of interbred transgenic mice expressing podocyte-restricted *APOL1-G0* or *APOL1-G2* simultaneously with a known disease trigger, HIV-1, using the Tg26 mouse model of HIVAN. Phenotypic comparisons revealed a protective effect on some glomerular pathology of the Tg26 kidney disease with co-expression of *APOL1-G0*. There was no difference with co-expression of *APOL1-G2* compared to the Tg26/HIVAN mouse model alone. *APOL1-G2* expression did not exacerbate disease but appears to have lost the protective functions exhibited by *APOL1-G0*.

## DISCUSSION

The mechanism by which the *APOL1* variants cause CKD remains unclear, and an unresolved issue is whether the *APOL1* variants are gain-of-function or loss-of-function mutations. There were no significant differences between the CKD phenotypes of the Tg26/*HIVAN4* model with co-expression of G2, indicating our prior report of an accelerated age-associated loss of podocytes in an unstressed state for the Tg-G2 mouse model was not reflecting a disease process that could be exacerbated with a stressor. On the contrary, the observed G0-dependent preservation of podocyte numbers and reduced glomerulosclerosis in the Tg26+G0 phenotype suggests G0 may be providing a mechanism to reduce stress-induced podocyte losses.

This preservation of podocytes may be related to podocyte hypertrophy, the only other significant difference between Tg26+G0 and Tg26+G2 mouse glomeruli. In glomerular disease, podocyte hypertrophy is a compensatory mechanism that maintains glomerular tuft coverage and preserves filtration barrier function in response to podocyte injury and loss.^31,32^ After podocyte loss, the remaining healthy podocytes hypertrophy to cover the vacant capillary surface. The observation that G0 mice had enhanced podocyte hypertrophy may suggest G0 function is involved in this compensatory stress response. The absence of hypertrophied podocytes in the Tg26+G2 mice may reflect either an absence of a hypertrophic response, or alternatively, the Tg26+G2 podocytes may have hypertrophied but then detached from the tuft and were lost. Studies of APOL1 function in *Drosophila* observed nephrocytes expressing G0 or G1 progressively hypertrophied and died as the fly aged, and this response was greater with G1 expression.^33^ If hypertrophy and cell loss is exaggerated with risk variant expression, additional hemodynamic factors^34^ or differences in cell-cell or cell-matrix attachment^35^ that occur in disease may contribute to the enhanced podocyte depletion.

Since APOL1 is only present in humans and a few other primates, transgenic mouse models are a pragmatic method to assess whole animal physiology of APOL1 function. However with any model system, there are limitations. The most significant concern is whether the human cellular pathways involving APOL1 function are present in mice. In addition, our mouse models restrict *APOL1* expression using the Nephrin promoter to podocytes and does not replicate the induction of *APOL1* expression by immune mediators and does not replicate expression in other sites, most notably expression in renal endothelium and in circulation. In our study, there was no significant effect on proteinuria, despite preservation of podocytes with reduced glomerulosclerosis. This may suggest additional pathogenic events in kidney cells other than podocytes may be important overall contributors to APOL1-associated CKD, which are not recreated in our mouse models of podocyte-restricted *APOL1* expression. Newly developed transgenic mouse models that express the entire *APOL1* gene including the flanking regulatory regions would be a better system to fully evaluate the stress-associated functions APOL1 in CKD.

This study is the first *in vivo* test of the function of kidney-expressed *APOL1* concurrent with a known human disease stressor. The function of *APOL1* in the presence of a disease stressor was only evident with G0, and not G2, indicating the *APOL1* risk variants may be loss-of-function mutations (summarized in **Figure 3**). Thus, the function of APOL1-G0 in the kidney may be to provide resilience to podocytes to tolerate disease stresses, and may explain why most individuals with a high risk genotype do not develop CKD. These studies would indicate the most logical approach for APOL1 targeted therapies is to restore *APOL1-G0* function in subjects with a high risk genotype.

## AUTHOR CONTRIBUTIONS

LAB, JFO, JRS, SM, PED, LB, DBT designed the study and interpreted data. ZW, LL, LAB, JFO, JRS conducted experiments. PED performed statistical analyses and prepared graphs. LB and DBT performed quantitative histopathology scoring. LAB wrote the manuscript and prepared figures. LAB, JFO, JRS, PED, LB, DBT edited the final manuscript. All authors reviewed and approved the manuscript for submission.

### ACKNOWLEDGMENTS

This work was supported by NIH grants DK097836 and DK108329. We thank the Vanderbilt Center for Kidney Disease for mouse renal function testing.

## DISCLOSURES

LAB, JFO, SM, JRS have received royalty payments for the commercial use of Tg-G0 and Tg-G2 transgenic mice.

