## Supplemental data for "*APOL1-G0* protects podocytes in a mouse model of HIV-associated nephropathy"

### Supplemental information

#### Figures

Supplemental Figure 1. Podocyte depletion occurs in the Tg26/*HIVAN4* mouse model.

Supplemental Figure 2. Podocyte density losses were not dependent on glomerular volume changes.

Supplemental Figure 3. No differences in survival rates of dual transgenics compared to Tg26/*HIVAN4*.

#### Tables

Supplemental Table 1. Histopathology scoring of Tg26/*HIVAN4* and *APOL1* dual transgenic mice.

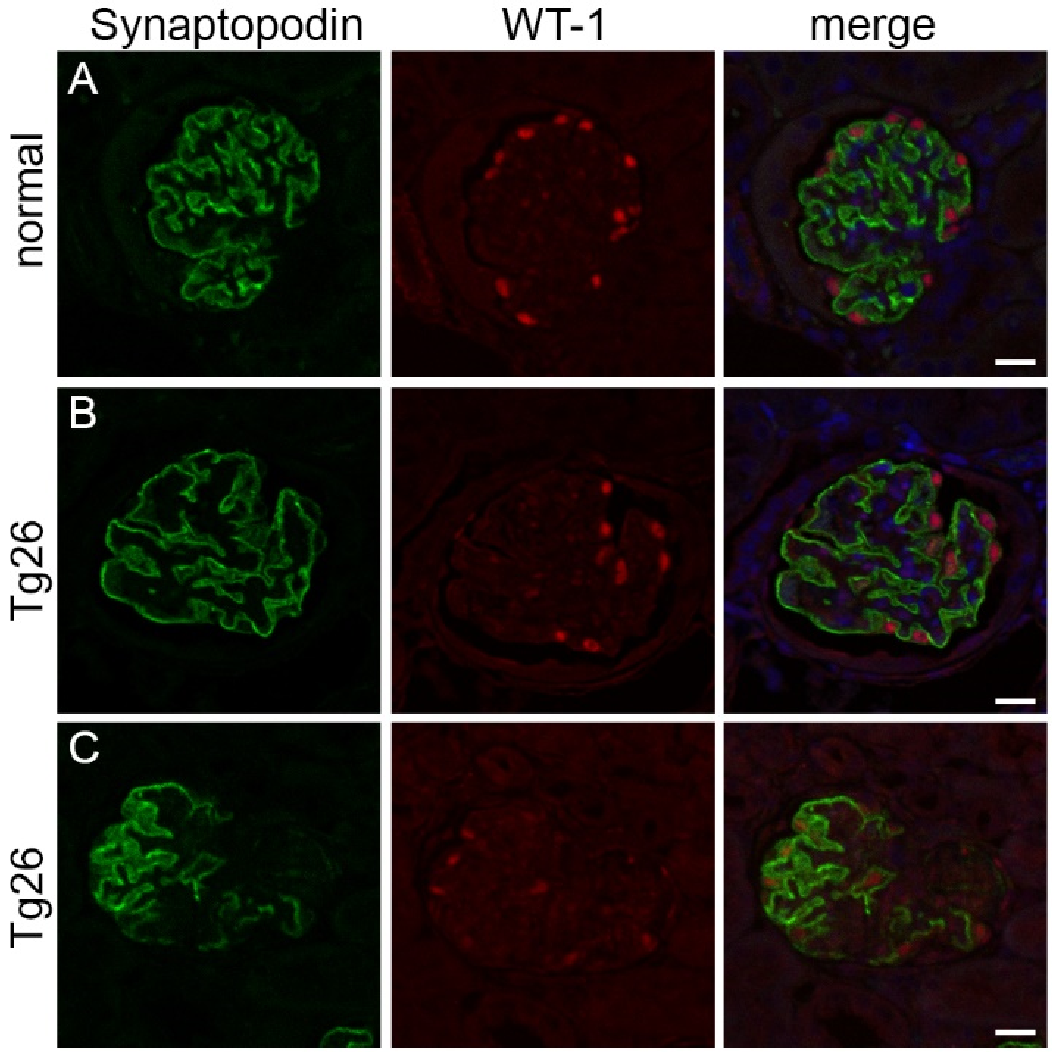


**Supplemental Figure 1. Podocyte depletion occurs in the Tg26/*HIVAN4* mouse model.** Immunofluorescence staining of WT-1 (nuclear) and Synaptopodin (cytoplasmic) in 200 day-old mouse glomeruli used to enumerate podocytes and estimate glomerular volumes respectively. **A**. Glomerulus from a normal mouse with typical distribution of podocytes and glomerular tuft structure. **B**. Glomerulus from a Tg26/*HIVAN4* mouse with a segmental loss of WT-1 positive nuclei, indicating loss of podocytes, but with uniform Synaptopodin positivity, indicating preservation of coverage of the tuft capillary surface (i.e. hypertrophy). **C**. Glomerulus from a Tg26/*HIVAN4* mouse with more advanced glomerular disease exhibiting both losses in number of WT-1 positive nuclei (and a qualitative reduced intensity of WT-1 staining), and segmental losses in Synaptopodin staining. Merge panel includes DAPI nuclear stain (blue). Scale bar=25µm.

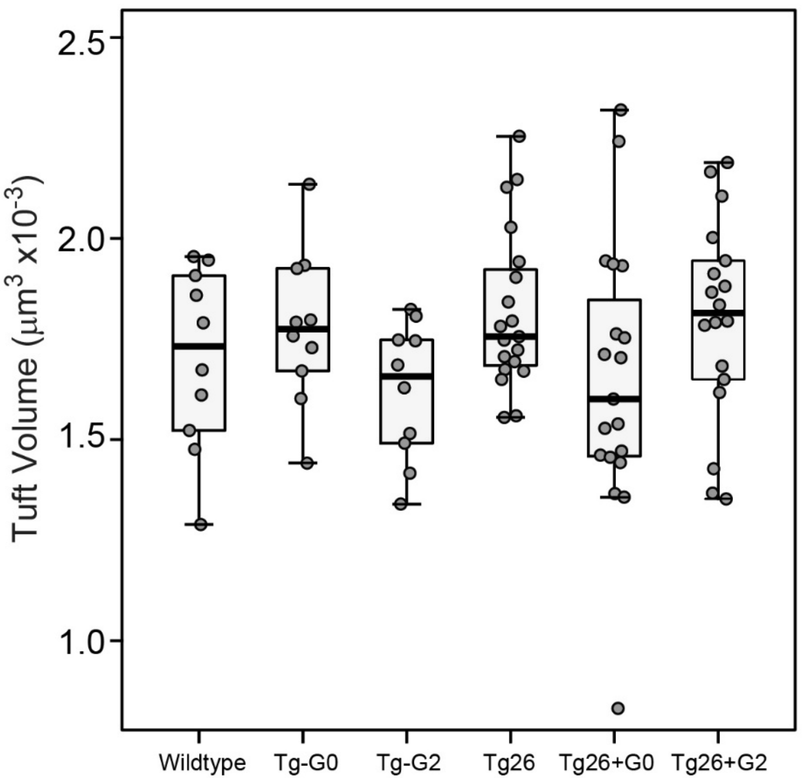


**Supplemental Figure 2. Podocyte density losses were not dependent on glomerular volume changes.** Glomerular tuft volumes were not significant differences between groups. Statistical comparisons were made to the relevant non-*APOL1* expressing group (Tg-G0 or Tg-G2 versus wildtype, and Tg26+G0 or Tg26+G2 versus Tg26). Each data point represents one mouse.

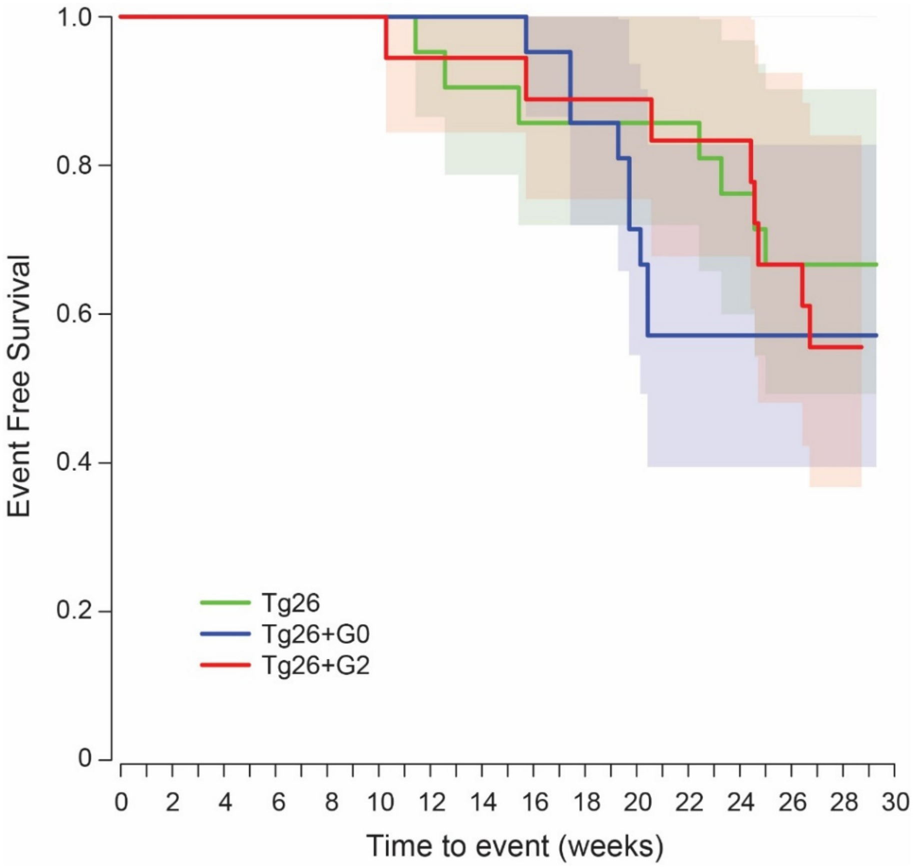


**Supplemental Figure 3. No differences in survival rates of dual transgenics compared to Tg26/*HIVAN4*.** Kaplan Meier plot of animals that died or reached one of the predetermined humane endpoints (this included renal failure or skin lesions that are typical of the Tg26 mouse model) prior to study endpoint of 200 days of age (29 weeks). Data are number of deaths (solid line) ± standard deviation (shaded area). There was no significant difference between groups.

**Supplemental Table 1:** Histopathology scoring of Tg26/*HIVAN4* and *APOL1* dual transgenic mice.

| | **Tg26 (n=21)** | ***P*** | **Tg26+G0 (n=21)** | ***P*** | **Tg26+G2 (n=18)** | ***P*** |
|---|---|---|---|---|---|---|
| # scored glomeruli per mouse (mean) | 201 | | 230 | | 210 | |
| Segmental sclerotic (% of total glomeruli) | 9.9% | ref | 6.6% | NS | 10.9% | NS |
| Global sclerotic (% of total glomeruli) | 3.7% | ref | 3.8% | NS | 5.2% | NS |
| Segmental collapse (% of total glomeruli) | 0% | ref | 0% | NS | 0% | NS |
| Global collapse (% of total glomeruli) | 0% | ref | 0.4% | NS | 0% | NS |
| Podocyte hyperplasia, any | 6.0% | ref | 6.0% | NS | 1.5% | NS |
| Podocyte hypertrophy, any | 10.0% | ref | 16.0% * | NS | 1.5% | NS |
| Tubular microcysts, mean±SD scored on 0-4 scale | 0.88 ± 1.27 | ref | 0.90 ± 1.47 | NS | 0.89 ± 1.33 | NS |
| Tubular atrophy, mean±SD scored on 0-4 scale | 0.07 ± 0.24 | ref | 0.05 ± 0.22 | NS | 0 ± 0 | NS |
| Interstitial fibrosis, mean±SD scored on 0-4 scale | 0.05 ± 0.22 | ref | 0.20 ± 0.52 | NS | 0.17 ± 0.51 | NS |
| Interstitial inflammation, mean±SD scored on 0-4 scale | 0.93 ± 1.29 | ref | 1.26 ± 1.66 | NS | 1.08 ± 1.41 | NS |

NS, not significant

* *P*=0.03 compared to Tg26+G2
